# Bee microbiomes in a changing climate: investigating the effects of temperature on solitary bee life history and health

**DOI:** 10.1101/2024.06.25.600696

**Authors:** Bailey L. Crowley, Robert N. Schaeffer

**Affiliations:** Department of Biology, Utah State University, Logan, UT, USA; Ecology Center, Utah State University, Logan, UT, USA

**Keywords:** *Osmia*, solitary bee, *Arsenophonus*, pollen provision, microbes, climate warming

## Abstract

Climate change is rapidly warming thermal environments, an important abiotic stimulus governing bee development and species interactions. Increasing evidence suggests that solitary bees rely upon pollen provision microbes for successful development. The effects of heat stress on provision microbiota and resulting consequences for larval health and development, however, remain to be examined. We performed an *in vitro* study to investigate the effects of thermal environment on provision microbiome composition and measured fitness outcomes for *Osmia lignaria* larvae. Elevated temperatures moderately increased the mean rank relative abundance of *Arsenophonus*, a putative son-killing symbiont. While pollen sterilization removed *Arsenophonus* from microbe-rich provisions, larval survivorship did not significantly differ between bees reared on microbe-rich and sterile diets. In contrast to previous research in solitary bees, larvae reared on sterile provisions weighed more and had higher total fat contents, with temperature moderating the degree of difference. As anticipated, we observed a negative relationship between the duration of larval development and temperature. Our results indicated that an intact provision microbiota may not always improve bee fitness, and that bee-microbe interactions during larval development may contribute to the size-shrinking effect observed for cavity-nesting bees under warming conditions.

## 1. Introduction

Globally, pollinators face a multitude of threats, including habitat loss, the introduction of invasive species, climate change, or a combination thereof [1]. Climate warming, in particular, can have both direct and indirect consequences for pollinators. For example, pollinating insects, such as bees, can be directly impacted by changes in temperature, an abiotic cue important for the timing of complex life history traits, such as development and emergence [2,3]. Shifts in life history traits may indirectly cause phenological mismatches between entomophilous plants and their pollinators, as these mutualists respond differently to changes in temperature [4]. In sum, bee responses to warming temperatures can have implications for population persistence, as well as broader effects on the species with which they interact.

An overwhelming majority of bee species are solitary, and they display considerable variation in functional traits, including their nesting preferences (ground vs. cavity) and overwintering stage (prepupae vs. fully pigmented adults). These functional traits influence how solitary bees are affected by variation in temperature [5]. For example, a recent study observed that the body size of cavity-nesting bees has declined more quickly than that of ground-nesting bees over the same 20-year period in response to rising temperatures [6]. Furthermore, recent experimental studies have found that cavity-nesting solitary bees experiencing elevated temperatures had increased fat body consumption, reduced longevity, and altered emergence phenology [7–11]. Remaining relatively unexamined is how temperature interacts with bee-associated microbiomes to affect host health.

Previous research has demonstrated that microbes associated with provisions (larval food) positively impact solitary bee health and development. For instance, a diet manipulation study found that, when fed sterile diets, solitary bee larvae grew slower, weighed less, and had reduced survivorship compared to their counterparts reared on microbe-rich provisions [12,13]. Upon further investigation, analyses comparing larvae reared on sterile and non-sterile provisions found more free fatty acids in bee in the latter treatment [14]. Although the mechanisms underlying these effects are unclear, early research suggests that microbes may permit larval access to bioavailable protein through degradation of recalcitrant pollen, or by serving as a dietary resource in and of themselves [15,16]. Thus, examining factors that may affect provision microbiome composition and function is essential to understanding drivers of bee health.

The activity of provision-associated microbes may be altered by abiotic factors, including temperature, which have potential downstream consequences for bee development and health. In some cases, increased temperature may be beneficial for the host through the indirect actions of symbionts. For example, high temperatures alter the symbiotic community of bumblebee gut microbes, resulting in reduced optimal growth temperatures for parasites and increased symbiont-pathogen competition [17]. However, destabilization of a symbiosis can also occur under thermal stress. For instance, the abundance of an obligate symbiont population was greatly reduced in a pest insect experiencing simulated warming and resulted in reduced fitness of the host [18]. While changes to the activities and functions of insect symbionts experiencing heat stress are still being uncovered, describing shifts in microbiome composition is a necessary step to elucidate these host-microbe interactions.

Here, we examined whether temperature shifts provision microbiome composition and the resulting effects on bee development and health outcomes using the solitary mason bee, *Osmia lignaria*. We implemented an *in vitro* diet manipulation experiment with a fully crossed 2 x 3 factorial design. The first factor, provision microbiome, consists of two levels - natural or sterile. Meanwhile, the second factor, temperature, includes three levels - cool, ambient, or warm. The simulated treatments emulated historic, current, and projected (2040-2099) temperatures for the Great Basin region (USA), where wild *O. lignaria* reside [19,20]. We used amplicon sequencing to describe shifts in provision microbiome composition and quantify bee fitness outcomes as larval body mass, total proportion of fat content, and development time (2nd to 5th instar). This experimental design allows us to examine how interactions between immature solitary bees and symbionts may have been altered during the past several decades of warming, and to predict how these relationships may change with continued increases in temperature.

## 2. Methods

### (a) Bees and *in vitro* rearing

*Osmia lignaria propinqua* (Megachilidae) is a native solitary mason bee found west of the Rocky Mountains (USA) [21]. *O. lignaria* offspring were collected from wild females nesting in artificial trap nests in Logan Canyon, Utah (UT), USA (41.89827°N, 111.57591°W) [22], located in the northeast corner of the Great Basin region. Here, bees have experienced between 0.3-0.6°C increase in temperatures over the past 100 years, and the western United States is expected to warm 2-5°C in the coming decades [19,20]. We regularly monitored the number of active and completed nests to determine peak nesting activity. Once peak nesting activity was reached, plugged nesting tubes were transported back to the lab and aseptically dissected in a laminar flow hood to reveal developing bees. For our diet manipulation study, we used a rearing protocol developed specifically for *O. lignaria* [23]. First, bees were grafted onto 1% agarose gels using sterile lab spatulas that were first submerged in sterile ringer’s solution [23]. We predominately grafted male *O. lignaria* because the species has a male-biased sex ratio. Male bees were selected from the first three brood cells (i.e., nearest the nest entrance), while females were only chosen from the last two brood cells that contained larger provisions [24]. We only grafted *O. lignaria* that were identified as eggs or first instar larvae, as bees in these early stages have not yet consumed any of the maternally allocated provision [25,26].

Provisions from randomly selected nests (*N* = 8) were subsampled to describe microbiome composition at the nest level. Then, provisions from all collected nests were homogenized, divided in half, and assigned to a microbiome treatment: microbe-rich or sterile. We homogenized provisions allocated to males and females together because there is no evidence of pollen microbiome composition between sexes [27]. To sterilize provision material, pollen was exposed to ethylene oxide (EO) gas overnight with one ampule of 17.5 g Anprolene in a benchtop fumigation chamber (Anderson Sterilizers Inc., Haw River, NC) [28]. Sterilized pollen was then rehydrated with sterile DI water to the initial weight prior to EO exposure. While it is unknown how this treatment affects the nutritional quality of the provision, previous research has demonstrated that bumblebees consume equal amounts of EO-treated and untreated pollen, suggesting that there is no effect on the palatability the pollen [28].

The following day, nesting tubes were used like cookie-cutters to weigh 190 ± 2 mg and 351 ± 2 mg of pollen for male and female bees, respectively. The amount of pollen allocated to immature bees was calculated as the average of 10 randomly selected provisions for each sex. These artificial brood cells were held upright in sterile 24-well plates to ensure bees do not fall off the provision, guaranteeing immobile larvae have access to food [23]. Male (*N* = 180) and female (*N* = 66) bees were randomized into treatments and grafted onto their respective provisions in the same manner as described above. To minimize the amount of handling time, all male bees were grafted within 24 h of nest dissection [23]. Immature females, however, were grafted 2 days after being removed from their natal nest. Bees were then placed into incubators (Percival I-30L, Geneva Scientific, Fontana, WI) with thermal environments approximating historic, current, and projected climates (hereafter cool, ambient, and warm) throughout the duration of their larval development. The ambient temperature treatment followed mean daily temperature fluctuations of June 2022 collected from data loggers (iButtonLink DS1925L-F5#, Whitewater, WI) located near the field site. Simulated cool and warm microclimates were programmed to be −1°C and +2°C relative to the ambient temperature treatment, respectively. Incubators across treatments were set at 60% relative humidity throughout the duration of the experiment. This experimental design produced a total of six different treatment combinations: cool-sterile (CN), cool-natural (CN), ambient-sterile (AS), ambient-natural (AN), warm-sterile (WS), and warm-natural (WN). Microbe-rich provisions without bees (351 ± 2 mg; *N* = 8 per treatment) were also placed within incubators to provide a baseline understanding of microbiome dynamics in response to temperature without potential modification by feeding larvae [29].

### (b) Bee response variables

Bees were monitored daily to track their development. Once bees reached the fifth instar, indicated by the production of frass [26], they were removed from their artificial brood cell and both the larva and provision weighed before being stored at −20°C until microbiome and lipid extractions were performed. To determine the total body fat content of fifth instar larvae, bees were first lyophilized and then submerged in 20-fold the volume of the tissue sample in 2:1 (v/v) chloroform-methanol solution. The chloroform-methanol solution was replaced every 24 h for two more subsequent days before larvae were once again lyophilized and reweighed. The proportion of total body fat was calculated as the difference between initial dry mass and dry mass following lipid extraction divided by the initial dry mass [11,30]. Of note, this method measures both fat bodies and non-storage lipids. However, in adult *O. lignaria*, this overestimation of storage lipids is negligible when using a methyl-chloroform extraction protocol [10,31].

### (c) DNA extractions

We prepared provisions for amplicon sequencing using DNA extraction protocols adapted from solitary bee microbiome papers [32–36] and incorporated advice for pollen metabarcoding [37,38]. First, samples were added to screw top microcentrifuge tubes containing about 100 µL of 0.1 mm glass beads, including reagent controls and 1 extraction blank for every 17 samples. To disrupt the cell walls of gram-positive bacteria, fungi, and pollen, samples were lysed with three cycles of 75 sec beating and 45 sec rest using a bead mill (Fisherbrand^TM^ Bead Mill 24 Homogenizer, Fisher Inc., Waltham, MA). Lysed samples were then processed using Qiagen Blood & Tissue kits with some slight adjustments: samples were treated overnight with 25 µL proteinase K (20 mg/mL) before centrifugation for 2 min at 6,000 rpm. Thereafter, kit instructions were followed.

We confirmed DNA concentration and the presence of microbial DNA in our samples using a Quant-iT High-Sensitivity dsDNA Assay kit (Invitrogen™, Thermo Fisher Scientific, Waltham, MA) and PCR, respectively. The bacterial 16S V4-V5 rRNA gene region was amplified using 5 μL Phusion™ Plus DNA polymerase (2x; Thermo Fisher Scientific, Waltham, MA), 0.75 μL of the forward primer 515B-F (5 μM; 5’-GTGYCAGCMGCCGCGGTAA-3’) [39], 0.75 μL of the reverse primer 926-R (5 μM; 5’-CCGYCAATTYMTTTRAGTTT-3’) [39], and 1.25 μL pPNA clamps (5 μM; PNA Bio, Newbury Park, CA) to prevent the amplification of plastid and mitochondrial DNA. To amplify DNA from bacteria, the PCR program included an initial denaturation step at 95°C for 3 min; 35 cycles of 95°C for 15 sec, 75°C for 10 sec, 50°C for 10 sec, and 72°C for 1 min; ending with a final extension step at 72°C for 10 min. PCR reactions to amplify the eukaryotic internal transcribed spacer 2 region (ITS2) of fungi included: 5 μL Phusion™ Plus DNA polymerase (2x), 0.5 μL of the forward primer ITS1-F (10 μM; 5’-TCCTCCGCTTATTGATATGC-3’) [40], and 0.5 μL of the reverse primer ITS4-R (10 μM; 5’-TCCTCCGCTTATTGATATCG-3’) [41]. The fungal PCR program began with an initial denaturation step at 95°C for 1 min; 35 cycles of 95°C for 1 min, 51°C for 1 min, 72°C for 1 min; followed by a final extension step at 72°C for 8 min. Samples that successfully amplified were submitted for amplicon sequencing.

### (d) Sequencing

Library preparation and Illumina MiSeq sequencing, producing 2 x 300 bp reads, was performed at Integrated Microbiome Resource at Dalhousie University (Nova Scotia, Canada). During sequencing, pPNA clamps and the same primers listed above were used, except the forward primer ITS86-F (5’-GTGAATCATCGAATCTTTGAA-3’) [42] was alternatively selected for the amplification of the ITS2 region. Samples containing DNA from bacteria was sequenced on a single run, while fungal samples were separated into two sequencing runs. The 16S and ITS2 raw reads were delivered as demultiplexed FASTQ files with adaptors already trimmed. Sequencing resulted in 794,418 total 16S reads with a mean ± standard error (SE) of 5,715.24 ± 1,145.43 reads per sample, while 428,074 total ITS2 reads were obtained with a mean ± SE of 2,218 ± 303.66 reads per sample. In R 4.3.1 [43], we inspected sequences using the DADA2 pipeline which filters and trims reads, dereplicates identical sequences, performs sample inference, joins paired end reads, and removes chimeric sequences [44]. Reads from the two fungal sequencing runs were separately passed through the DADA2 pipeline before being merged into a single dataset.

To ensure quality reads, bacterial sequences were trimmed at 250 bp and 200 bp for the forward and reverse reads, respectively, where the quality scores dropped below thirty. Taxonomy of bacteria was assigned using the 16S rRNA Ribosomal Database Project v18 naïve Bayesian classifier [45]. We removed sequences classified as cyanobacteria, chloroplast, mitochondria, Archaea, and Eukarya from the bacterial dataset. ITS2 primers were removed from raw reads using cutadapt v4.6[46], and we filtered ASVs that were not identified to at least family. We used the UNITE ITS v9.0 database for taxonomic assignment of fungi [47]. Using the *decontam* package, contaminant reads were removed from samples based on prevalence in reagent and extraction blanks [48]. After controlling the quality of sequences, we retained a total of 1,930 paired-end reads from 138 ASVs in our bacterial dataset containing 30 samples. In the fungal dataset, we preserved 10,478 paired-end reads from 266 ASVs in our 68 samples. Mean reads per sample were 64.33 (± 7.35 SE) and 154.09 (± 21.25 SE) for bacteria and fungi, respectively. Albeit low, our rarefaction curves suggest that extraction methods sufficiently captured the diversity of the provision microbiome (Figure S1). While other studies find higher read counts per provision in solitary bee species [32–36], the age of pollen is typically not reported. We sample provisions when larvae were fifth instars; the viability of microbial taxa in aged solitary bee pollen has yet to be examined. In fact, the microbiota of honey bee stored pollen undergoes drastic reductions in diversity and culturable microbes, ultimately becoming dominated by osmotolerant bacteria [49].

### (e) Statistical analyses

All statistical analyses were performed in R 4.3.1 [43]. Health and life history data from female bees were excluded from analyses because they died rapidly, likely because of delayed grafting – females were off their provisions for 48 h (BLC personal observation). We implemented linear mixed effects (LME) models using the lme4 package to assess the effectiveness of simulated climatic treatments and to analyse the effects of treatments on bee response variables [50]. In the models assessing differences in mean temperature and relative humidity, we used day of year as a random intercept to account for multiple daily temperature readings. Meanwhile, grafting stage (egg or 1st instar) was used as a random intercept in models testing the effects of microbiome and temperature treatments on the biomass, proportion of fat content, and development time of male larvae. Residuals for larval development time were not normally distributed; thus, this response variable was analysed using a generalized mixed effects model (GLMM) with a gamma error distribution. We estimated the *p*-values of linear mixed effects models using the Kenward-Rogers approximation. We used the emmeans package to examine pairwise comparisons, adjusting *p*-values with the Tukey’s Honest Significant Difference method (Tukey’s HSD) [51]. Prior to analysing survivorship data, we removed male bees that died within 48 h of grafting, which we attribute to physical damage inflicted during handling. Log rank and Gehan-Breslow generalized Wilcoxon tests were used to compare median survival time of male bees across treatments using the survival package [52]. In addition, we used the survreg function to fit an accelerated failure time (AFT) model to determine the effects of temperature and microbiome treatments on survival time, including grafting stage as a fixed effect.

We qualitatively describe the microbiomes of provision samples that were not consumed by larvae but exposed to temperature treatments given the small number of these samples. Provisions containing developing bees (males and females) were statistically analysed. We removed the only cool-sterile sample from our bacterial dataset prior to statistical analysis because it contained two reads. Alpha diversity measures, including Shannon’s Diversity Indices and observed richness, were calculated using the estimate_richness function from the phyloseq package [53]. We assessed the effects of bee sex and temperature treatment on alpha diversity using a linear mixed effects model (LME) with the nlme package, including grafting stage as a random effect [54]. In addition, we quantified Pielou’s evenness to examine the effect of bee sex, thermal environment and microbiome treatments on the distribution of microbial ASVs amongst our treatments. We also performed a Kruskal-Wallis test *a posteriori* to evaluate shifts in mean relative abundance of the dominant bacterial genus, *Arsenophonus*, in response to our climatic treatments.

To test the effects of microbiome treatment and thermal environment on the composition of bacterial and fungal communities, we performed permutational multivariate analyses (PERMANOVA 999 permutations) using the adonis2 function from the vegan package [55]. Beta diversity analyses were applied to Bray-Curtis distance matrices generated from relative abundance datasets. When an overall significant effect was found, pairwise comparisons with Benjamini-Hochberg (BH) correction of *p*-values were performed using the RVAideMemoire package [56]. The PERMANOVA assumption of homogenous group dispersion was inspected with the betadisper function. When significant, we followed up with pairwise comparisons using the Tukey’s HSD method. Differences in microbiome composition between treatments were visualized with principal coordinates analysis (PCoA) plots.

## 3. Results

### (a) Effectiveness of climatic and microbiome treatments

The mean thermal environment in the warming treatment was 2.2°C higher than the ambient treatment; meanwhile, the mean temperature of the cool treatment was 0.8°C below the mean ambient temperature (LME, *F*_2,10227_ = 117.65, *p* < 0.001; figure 1). The maximum temperatures bees experienced in each microclimate were 23.2°C, 25°C, and 27.2°C for the cool, ambient, and warm treatments, respectively. These climatic manipulations approximate historic (*c*. 1900s), current, and conservative projected (*c*. 2050-2100) mean daily temperatures for the Great Basin region [19,20]. Although relative humidity was programmed to remain the same across incubators regardless of simulated microclimate, this variable significantly differed between thermal environments (LME, *F*_2,11183_ = 343.15, *p* < 0.001; figure 1). However, mean relative humidity was 61.74%, 60.60%, and 61.33% for the cool, ambient, and warm microclimates, respectively. These minimal differences in relative humidity between treatments likely did not affect bee fitness outcomes and shifts in microbiome composition.

**Figure 1.**
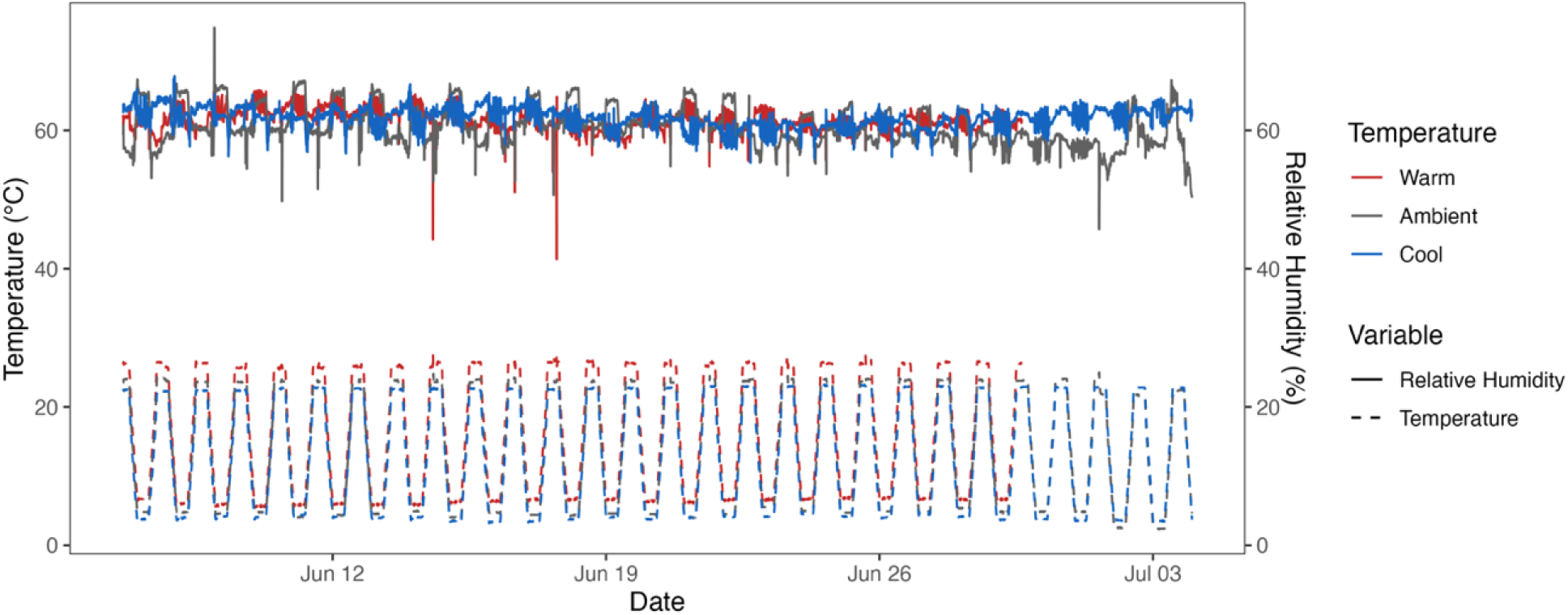
Simulated temperature treatments emulating past (*c*. 1900), current, and projected (*c.* 2050-2099) temperatures for the Great Basin region. Mean temperature for the cool, ambient, and warm treatments were 12.9°C, 13.7°C, and 15.9°C, respectively. Relative humidity was approximately 60% for all treatments across the length of the experiment. Temperature and humidity loggers were stopped once all bees within the treatment had reached the fifth instar.

Nest-level sampling of pollen prior to homogenization revealed that the bacterial genus assigned the most read counts, *Arsenophonus*, did not appear consistently in natal nests; however, after mixing, our mass of pollen contained this genus as a dominant member of the provision microbiota (figure S2). The fungus, *Cladosporium*, was found in nearly all nest-level samples of provisions, while other genera appear more sporadically and in low relative abundances (figure S2). Inspection of our pollen mass after homogenization suggests that we sufficiently mixed provisions prior to reallocating larval food to *O. lignaria* offspring, as *Cladosporium* remained the most dominant fungal genus and other taxa were found in low relative abundances (figure S2).

Pollen sterilization was effective at eliminating bacteria, as amplicon sequencing did not detect reads from this kingdom in EO-treated pollen. Fungi were still detected in sterilized pollen, but analyses indicated that our method reduced species richness (LME, *F*_1, 38_ = 9.90, *p* = 0.003) and had significant effects on provision composition (PERMANOVA, *R^2^*= 0.12, *F* = 5.76, *p* = 0.001). Upon examining Pielou’s evenness, we found a marginal effect of temperature on the distribution of bacterial taxa in microbe-rich provisions (LME, *F*_1,11_ = 3.58, *p* = 0.09; figure S3), suggesting that thermal environment may affect the relative abundance of certain genera. In contrast, temperature did not have an effect on the evenness of fungal taxa (LME, *F*_2, 31_ = 0.33, *p* = 0.72; figure S3) but pollen sterilization had a marginal effect (LME, *F*_1,31_ = 3.55, *p* = 0.07; figure S3).

### (b) Overall microbiome composition

Most of the taxa present in provisions were represented by common soil-borne microbes and plant pathogens, such as Bacillaceae, Bradyrhizobiaceae, Comamonadaceae, Cladosporiaceae, Valsaceae, and Pucciniaceae. In addition, two insect-associated genera were present: *Arsenophonus* and *Ascosphaera*, accounting for 43.26% and 0.07% of total bacterial and fungal reads, respectively. In nearly all provision samples with feeding larvae present, *Arsenophonus* and *Cladosporium* were found in high relative abundance (figure 2, figure S4). Provisions without feeding larvae had lower mean bacterial read counts (36 ± 12.32 SE) compared to those with actively feeding larvae (82 ± 21.16 SE). Similar patterns were also observed with fungi: provisions had higher fungal read counts when larvae were present (205.68 ± 63.49 SE) than absent (56.82 ± 20.09 SE). Nearly all bacterial and fungal genera detected in pollen without actively feeding larvae were also found in provisions containing developing bees (figure S4, figure S5). Indeed, overall bacterial (PERMANOVA, *R^2^* = 0.04, *F* = 1.00, *p* = 0.42) and fungal (PERMANOVA, *R^2^* = 0.02, *F* = 0.99, *p* = 0.41) composition in provisions did not differ depending on the presence or absence of larvae. However, provisions with actively feeding larvae contained more bacterial and fungal genera that were not found in pollen without bees, albeit in low relative abundances (figure S4, figure S5).

**Figure 2.**
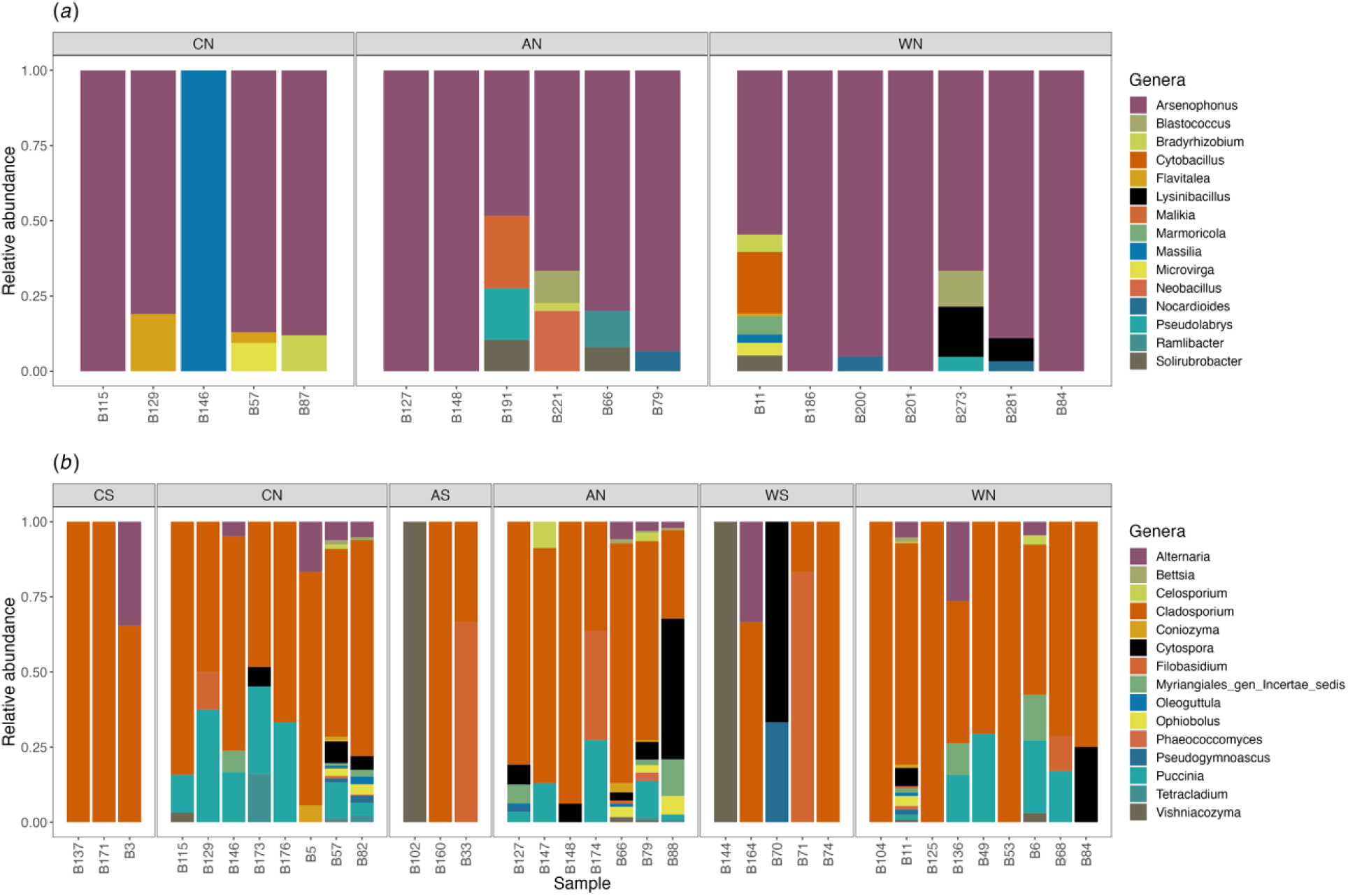
Relative abundance (proportion of sequence reads) of the top 15 (*a*) bacterial and (*b*) fungal genera present in *Osmia lignaria* provisions with male and female bees.

### (c) Impact of thermal environment on microbiome diversity and composition in provisions with actively feeding larvae

Thermal environment had no effect on the alpha diversity of bacteria, quantified as Shannon’s Diversity Index (LME, *F*_2,11_ = 0.24, *p* = 0.79; figure S6) and observed richness (LME, *F*_2,11_ = 0.43, *p* = 0.66; figure S7). Although we did not find a significant effect of temperature on the mean relative abundance of *Arsenophonus* (Kruskal-Wallis, *H*_2_ = 0.78, *p* = 0.68), the warm treatment trended toward a higher mean (0.81 ± 0.07 SE) than both the cool (0.78 ± 0.06 SE) and ambient (0.68 ± 0.10 SE) microclimates (figure 3). Thermal environment did not have an effect on the observed richness of fungal ASVs across treatments (LME, *F*_2,38_ = 0.88, *p* = 0.42; figure S6), but a marginal effect was found for Shannon’s Diversity Index (LME, *F*_2,38_ = 2.82, *p* = 0.072; figure S5). Bee sex was a significant predictor in the model of the observed richness of bacteria (LME, *F*_1,11_ = 5.63, *p* = 0.037), however there were no significant pairwise comparisons. Likewise, bee sex was a significant covariate in the model of fungal Shannon’s diversity (LME, *F*_1,38_ = 6.26, *p* = 0.016), but the only significant pairwise comparison detected were between sterile and natural provisions.

**Figure 3.**
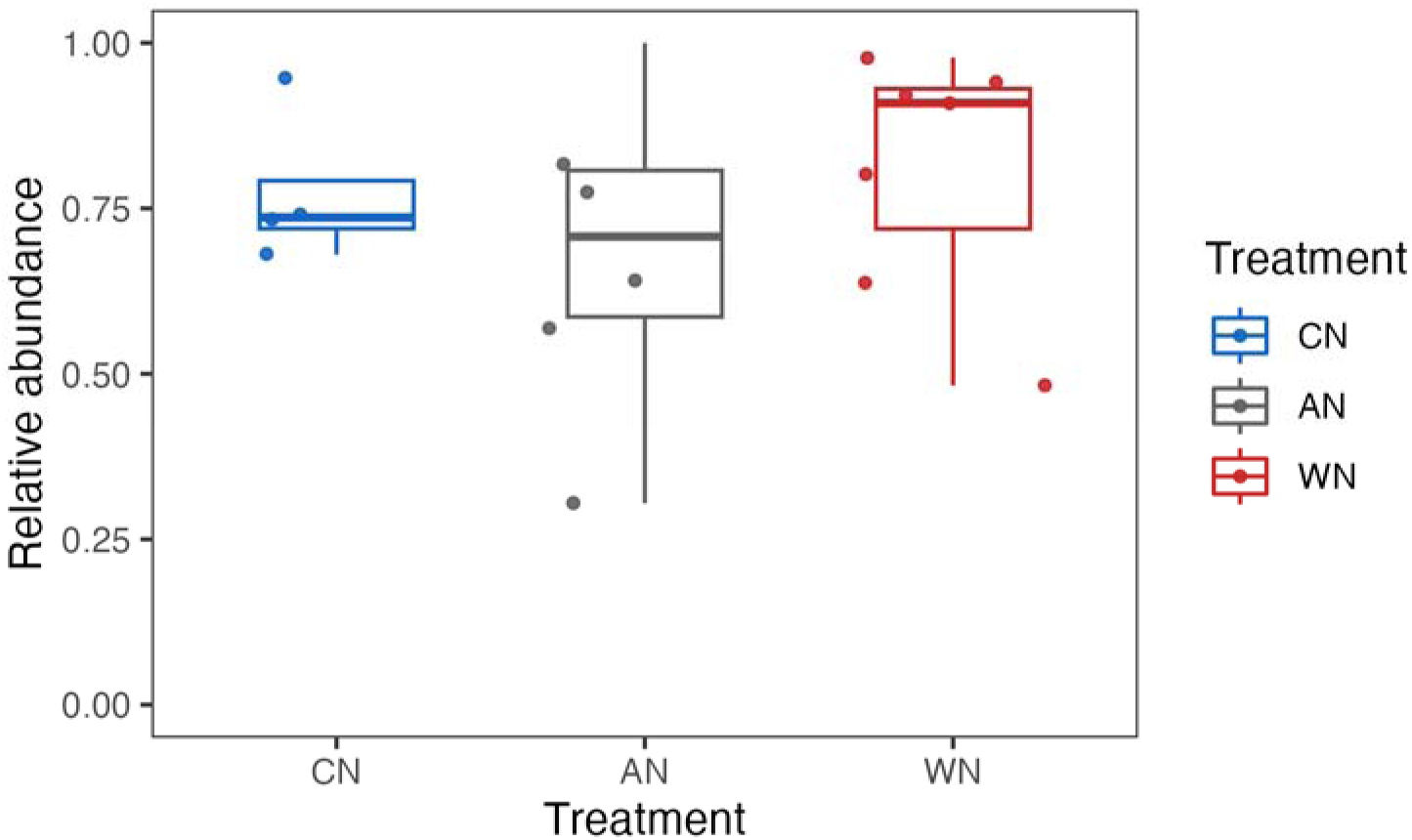
Mean rank relative abundance of *Arsenophonus* in provisions with bees. Kruskal-Wallis tests did not indicate any significant differences between treatments due to temperature treatment nor sex.

Both temperature (PERMANOVA, *R^2^* = 0.13, *F* = 0.126, *p* = 0.086) and bee sex (PERMANOVA, *R^2^* = 0.07, *F* = 1.38, *p* = 0.052) had a marginal effect on the composition of bacterial ASVs in provisions with actively feeding larvae. Dispersion between groups was not violated due to temperature treatment (ANOVA, *F* = 0.02, *p* = 0.98) nor bee sex (ANOVA, *F* = 0.19, *p* = 0.66). Temperature (PERMANOVA, *R^2^* = 0.04, *F* = 0.97, *p* = 0.47) and bee sex (PERMANOVA, *R^2^* = 0.02, *F* = 1.19, *p* = 0.29) did not have a significant effect on fungal composition. In assessing the PERMANOVA assumption of fungal composition, homogeneous group dispersion was not violated in pairwise comparisons between microbe-rich provisions. Upon visual inspection in ordination space, the provision microbiome did not cluster by temperature treatment (figure S8).

### (d) Larval health and development outcomes

Microbiome treatment had an overall effect (LME, *F*_1,53.11_ = 8.02, *p* = 0.007) on the biomass of male fifth instar larvae. Pairwise comparisons revealed a moderate effect (Tukey’s HSD *p* < 0.08) when comparing sterile and microbe-rich provisions within each temperature treatment as well as between WN versus both CS and AS. While the thermal environment did not have an overall effect on larval body weight (LME, *F*_2, 53.22_ = 1.28, *p* = 0.29), the differences in mean biomass between larvae reared on sterile versus microbe-rich provisions was incrementally greater with increasing temperatures. Specifically, bees reared on sterile pollen were 3.88%, 22.3%, and 28.7% larger on average than larvae consuming microbe-rich provisions in the cool, ambient, and warm microclimates respectively (figure 4*a*).

**Figure 4.**
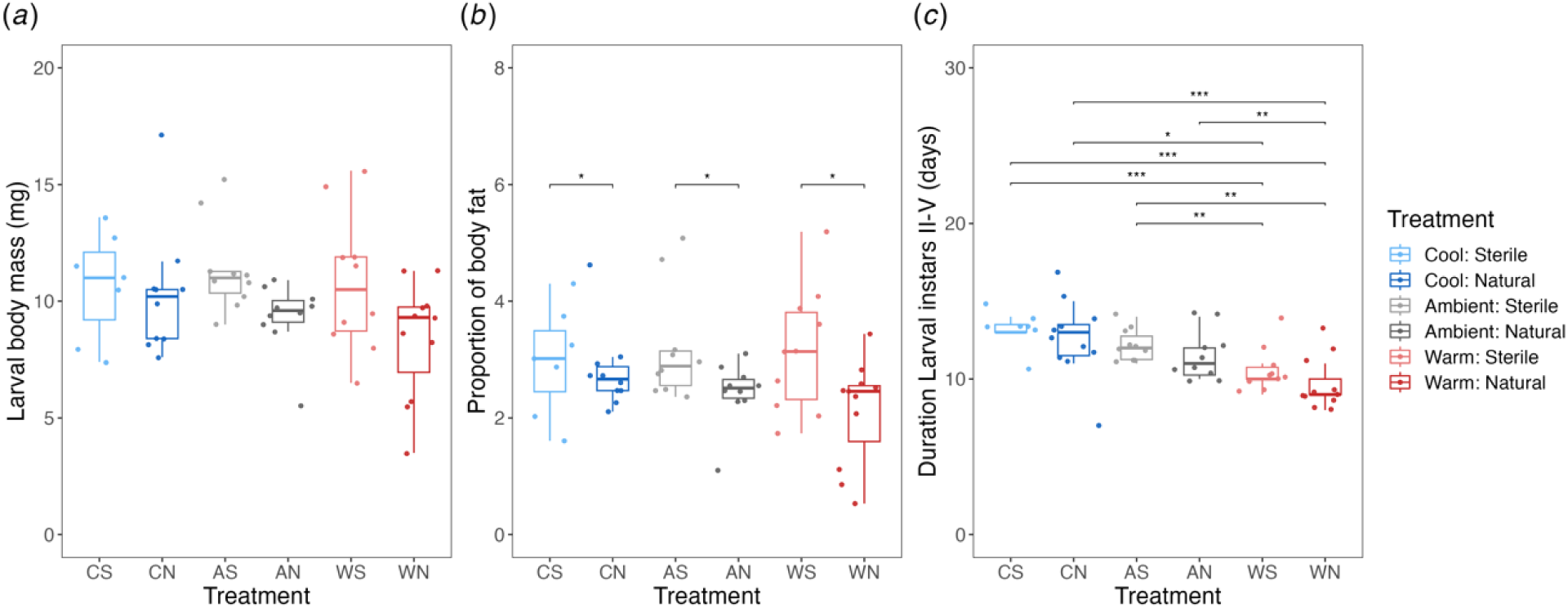
Male bee health and life history outcomes, including (*a*) body mass, (*b*) proportion of body fat, and (*c*) development time from the second to fifth larvae in response to temperature and microbiome manipulations. Fifth instar larvae were used to determine body mass and the proportion of body fat. Asterisks represent significant differences, where ‘*’ indicates a *p*-value < 0.05, ‘**’ *p* < 0.01, and ‘***’ *p* < 0.001.

Similarly, the presence or absence of microbes in the larval diet had a significant effect on the proportion of total fat content in male immature bees (LME, *F*_1,53.10_ = 10.42, *p* = 0.002). Thermal environment did not have an overall effect (LME, *F*_2,53.19_ = 0.82, *p* = 0.44) on the fat content of fifth instar larvae; however, increasing temperatures exacerbated the differences in mean total fat content between bees reared on sterile and non-sterile provisions. In particular, larvae reared on sterile provisions had 6.45%, 30.7%, and 49.8% higher total fat content on average than bees fed microbe-rich provisions (figure 4*b*).

The duration of larval development was significantly affected by nest temperatures (GLMM, *X^2^* = 20.08, *p* < 0.001) but was unaffected by microbiome treatment (GLMM, *X^1^* = 2.58, *p* = 0.11). Further investigation revealed significant differences in all pairwise comparisons between larvae reared in the warm nest environment compared to those in the cool and ambient microclimates, irrespective of microbiome treatment (figure 4*c*). Median survival time was not significantly different between treatments (log rank test: 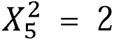, *p* = 0.8); Gehan-Breslow-Wilcoxon test: 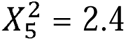, *p* = 0.8) (figure 5). These findings are supported by the AFT model (*X*^5^ = 4.78, *p* = 0.44): neither temperature (*p* = 0.40) nor microbiome (*p* = 0.66) treatments had a significant effect on larval survivorship.

**Figure 5.**
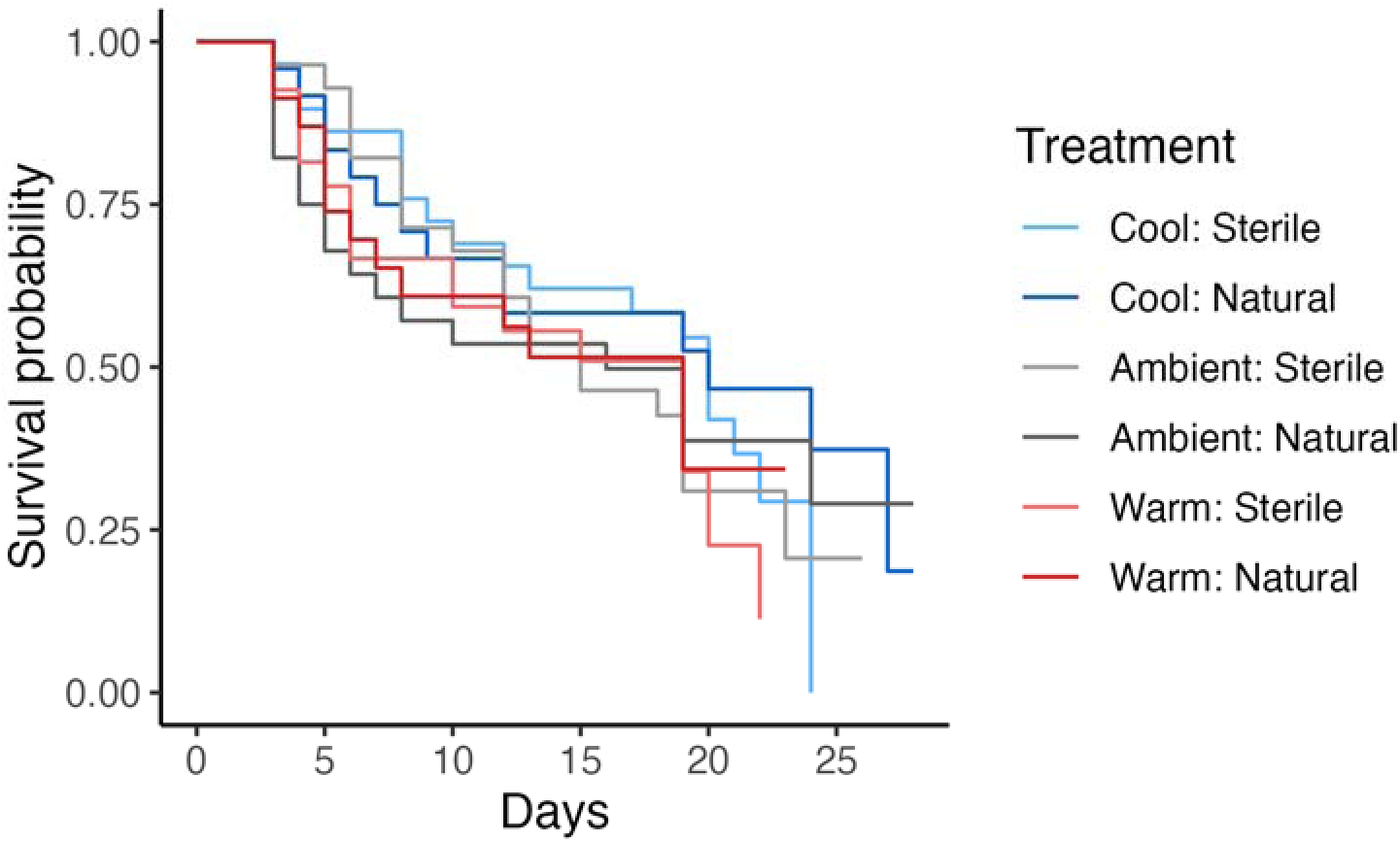
Kaplan-Meier survival plot of male *Osmia lignaria* larvae across treatments. Bees that died within 48 h of grafting were excluded. An accelerated failure time model did not reveal significant differences in survivorship across treatments due to microbiome nor temperature treatment.

## 4. Discussion

Solitary bees are thought to derive benefits from the provision microbiota, either as dietary resources or by pre-digesting pollen [12–14,16]. Differences in thermal optima between larvae and symbionts have unknown consequences for developing solitary bees. We performed an *in vitro* study to investigate the effects of temperature on provision microbiome structure and determined the outcomes for the health and development of male *O. lignaria* larvae. We found trends that point to temperature-mediated effects on the mean relative abundance of *Arsenophonus*, a putative son-killing bacterium in solitary bees. In contrast to expectations, we did not observe differences in survivorship between larvae reared on microbe-rich and sterile diets, although pollen sterilization was effective at eliminating *Arsenophonus*. Upon examining the effects of microbiome treatment on larval biomass and fat content, we found that larvae reared on sterile provisions weighed more and had higher total fat contents. Within each thermal environment, the differences in mean biomass and fat content between bees reared on sterile versus microbe-rich provisions increased with warming temperatures. We also found a negative relationship between temperature and the duration of larval development. Our data presents evidence that the maternally provided provision microbiota may not be beneficial to bee health in all circumstances. In addition, this study highlights that bee-microbe interactions may be mediated by temperature, with potential consequences for larval fitness such as the decrease in bee body size observed over the past several decades of warming climates [6,57].

While most microbial taxa were not specific to *O. lignaria*, the most dominant bacterial genus detected, *Arsenophonus*, is a known symbiont of plants and insects. The *Arsenophonus* genus is monophyletic group contains symbionts which exhibit exceptionally diverse phenotypes, including phytopathogenicity, obligate endosymbiosis, and reproductive parasitism [58–60]. The 16 ASVs assigned to *Arsenophonus* in this study had on average 98.92% sequence similarity to the son-killer, *A. nasoniae* (table S1) [61–63]. Surveys of social and non-social bees have detected *Arsenophonus* in pollen, bee body surfaces, crops, and midguts [64–72]. The *A. apicola* strains isolated from honey bees have also been implicated as a male-killing symbiont [73,74]. Although *A. nasoniae* originally described in wasps lacks genes for the uptake of glucose and metabolisms of key amino acids [75], *A. apicola* has the ability to utilize more carbon sources [74], likely the reason why the bee associate can be found in flowers and stored pollen. Laboratory studies discovered that *Arsenophonus* is transmitted from mother to offspring via inoculation of the egg surface, eventually localizing in the midgut of infected individuals and then may cross the gut epithelium [70,76,77]. In a recent study that characterized the microbiome of *O. lignaria* across bee development, *Arsenophonus* was found in low relative abundance in early developmental stages and only reached higher read counts in the guts of adults [71], suggesting that similar infection processes may occur in solitary bees.

In the present study, some *O. lignaria* larvae successfully developed in the presence of the putative son-killing bacterium. While investigating the transmission of *Arsenophonus* between honey bee nestmates, Drew et al. (2021) documented that a subset of adult bees lost the infection over the course of the experiment, likely by shedding the parasite in faeces [70]. We speculate that the surviving bees in our study voided *Arsenophonus* in their digestive tract once they reached the fifth instar stage, or shed the putative reproductive parasite while moulting. When tracking *A. nasoniae* infection processes *in vivo*, the parasite was found on the host digestive tract, ovipositor, and exuviae of infected male wasps [78], suggesting that this bacterium has the ability to metabolize chitin. Indeed, genes for the production of a novel chitinase and chitin-binding proteins have described in *A. nasoniae* and are likely responsible for the ability of the parasite to occupy this niche [75].

Although our sterilization method eliminated *Arsenophonus* and other bacteria from microbe-deficient pollen, larvae reared on sterile provisions had comparable survivorship to bees consuming microbe-rich diets. These results contradict previous research strongly suggesting that larval survivorship depends on the presence of microbes in larval food [12–14]. Preliminary data from a recent *in vitro* study, however, also indicates that provision microbes can have adverse consequences for solitary bee fitness: the addition of *Apilactobacillus micheneri* (previously *Lactobacillus*) in the diet of megachilid larvae increased mortality [79], an unexpected result given the frequent detection in solitary bees [66,80,81] and known functions of lactobacilli in eusocial bees (e.g., preventing the spoilage of stored pollen and defense against pathogens) [82,83]. Furthermore, Anderson et al. (2014) used microscopy and culture-dependent methods to challenge the long-held belief that microbes in the stored bee bread of honey bees aid in the pre-digestion pollen: upon visual inspection, microbes were greatly outnumbered by uncompromised pollen grains, and counts of viable microbes markedly fell after 96 h of storage [49]. Collectively, these findings suggest that the maternally allocated provision microbiome may not always benefit larval health. Future research should leverage experimental studies to investigate the functions of exosymbionts found within solitary bee provisions.

We also found contradictory evidence regarding the importance of the provision microbiota for larval health: immature bees reared on pollen without microbes weighed more than those feeding on microbe-rich provisions. Past studies found that larvae acquire fatty and amino acids from microbial prey within provisions, resulting in heavier bees than those reared on sterile pollen [12–14]. These opposing results may be explained by differences in pollen sterilization methods: in choice assays, EO exposure does not affect the palatability of pollen to bumblebees [28], but it is unknown whether soaking provisions in ethanol [12–14] affects pollen consumption in solitary bees, perhaps leading to differences in biomass, fat content, and trophic positions. Of note, while not statistically significant, bumblebees consumed less pollen per day when fed EO exposed pollen compared to the no treatment control [28]. This likely contributed to slight differences in biomass and fat content between bees reared on sterile versus microbe-rich provisions, though most pollen consumption occurs while fifth instar larvae in *Osmia* spp. Indeed, *O. lignaria* body size is correlated with the weight of the provision [84].

In this study, *O. lignaria* larvae feeding on microbe-depleted provisions had higher total fat content than those reared on microbe-rich provisions. *Arsenophonus* has been detected in the fat bodies of diverse insect taxa, including wasp larvae [78], adult planthoppers [85], and bumblebees co-infected with *Apicystis bombi* [86], a parasitic protist clearly documented to cause fat body depletion [87]. Although *Arsenophonus* has been detected in the fat bodies of these diverse insect taxa, the capacity of this bacterium to metabolize host lipids has not been studied, rather most research on reproductive parasites have focused on their ability to manipulate the sex ratio of their hosts. Thus, whether *Arsenophonus* is responsible for the increased fat body consumption in larvae reared on microbe-rich provisions remains to be determined. We also found that *O. lignaria* larvae reared on microbe-rich provisions in the warmest microclimate had the largest difference in mean biomass and fat content compared to their counterparts reared on microbe-depleted provisions. Thus, *Arsenophonus*-mediated lipid consumption may be affected by environmental temperatures. Alternatively, heat-stress may affect the nutritional quality of pollen: although provision mass was comparable between treatments*, O. lignaria* larvae consuming provisions with pollen collected from plants that experienced a heatwave had lower survivorship than immature bees fed pollen from non-stressed plants [88]. Regardless of the mechanism, reductions in total fat content of *O. lignaria* larvae reared on microbe-rich provisions, as found in this study, is concerning, as lipids obtained early in bee development are converted into fat reserves used during diapause. In overwintering adults, warmer temperatures elevated bee metabolism and lead to increased fat body depletion [7,9,10]; these effects may be intensified if larvae do not acquire enough fat storage prior to pupation. Furthermore, emerged adults with reduced fat stores have shorter lifespans and vigor [7–9], which may result in negative consequences for the reproductive success of female bees by impacting ovarian development [89].

Most research studying the effects of temperature on development in solitary bees have focused primarily on the mature stages given the importance of timing emergence phenology with flowering periods and adult vigor in managed species. However, bees have stage-specific temperature requirements. For instance, *O. cornifrons* has a narrow thermal window for successful development of immature bees: eggs only remained viable between 13-29°C and larvae were sensitive to temperatures outside of 21-29°C [90]. The consequences of a warmer nest environment on larval survivorship depends on the frequency and intensity of thermal stress. For example, exposure to consistently elevated temperatures accelerated larval development [91] but brief exposure to heatwaves slowed growth in *Osmia* spp. [92], suggesting that temperatures above a certain thermal tolerance can inflict physiological damage that affects the uptake of food by immature bees. Our treatments fell within the thermal window of *Osmia* spp. [89], and thus were not likely the cause of the high larval mortality observed in this study. We found that increases in nest temperatures shortened the duration of *O. lignaria* larval development. Other studies have shown that warming also shortens the length of development from the pupal to adult stages in *Osmia* spp. [93]. Entering the adult stage early and continuing to experience elevated temperatures while overwintering has adverse consequences for bee fitness, including reduced biomass and increased mortality [7–10, 93]. In addition, warming can have non-linear effects on nest emergence – either advancing [8,94] or delaying [11] emergence. Shifts in bee emergence can lead to phenological mismatches with floral resources [95], which may be especially detrimental for early-spring bees, such as *Osmia* spp.

Here, we found that the duration of larval development shortened with warming temperatures. Simulating a warmer climate increased the mean relative abundance of the putative son-killer *Arsenophonus*, yet larval survivorship did not differ between bees reared on microbe-rich versus sterile provisions. By comparing the difference in larval biomass and fat content between bees reared on pollen with and without microbes, we provide evidence that an intact provision microbiota may not always be beneficial; however, the degree by which larval biomass and fat content differed was moderated by temperature. These results suggest that provision exosymbionts may have contributed to the decrease in body size observed during the past several decades of warming.

## Data accessibility

Additional data are available as electronic supplementary material. The datasets, metadata, and R scripts generated and/or analyzed during the current study are openly available in the GitHub repository, https://github.com/BaileyLC/Osmia_climate. Raw sequence data were deposited in the NCBI Sequence Read Archive (SRA) under the BioProject numbers PRJNA1089148 and PRJNA1089149 with their respective accession numbers.

## Authors’ contributions

B.L.C and R.N.S conceived the study and authored drafts of the article; B.L.C. performed the experiments and data analyses.

## Acknowledgements.

The authors thank Drs. Karen Kapheim and Carol von Dohlen for their feedback on an early draft of this manuscript. We also thank the USDA-ARS Pollinating Insect-Biology, Management, Systematics Research facility for donating materials required to trap wild mason bees and sharing protocols for the rearing of *Osmia lignaria*.

---

**Figure S1.**
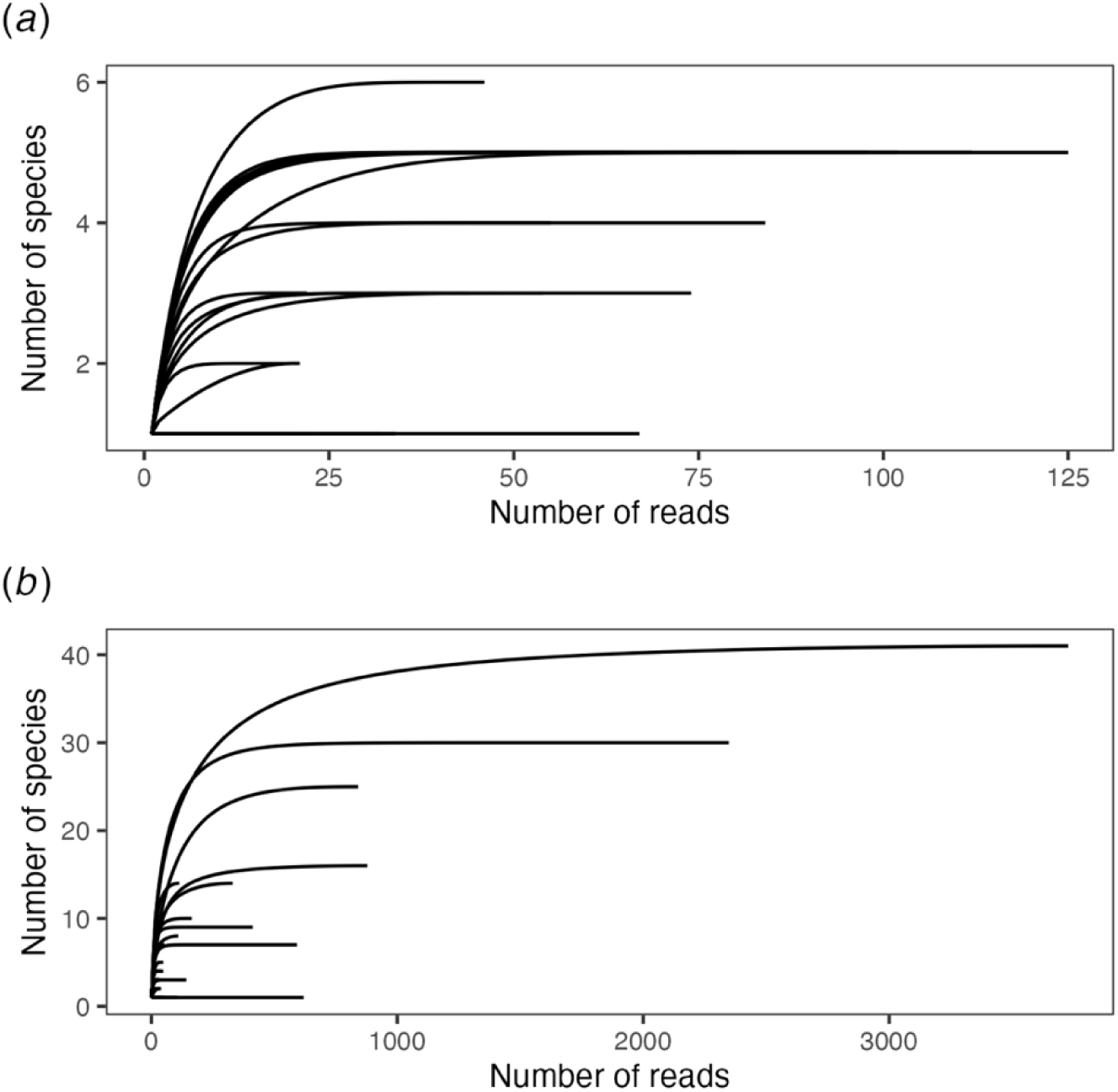
Rarefaction curves showing observed (*a*) bacterial and (*b*) fungal species richness in provisions with bees.

**Figure S2.**
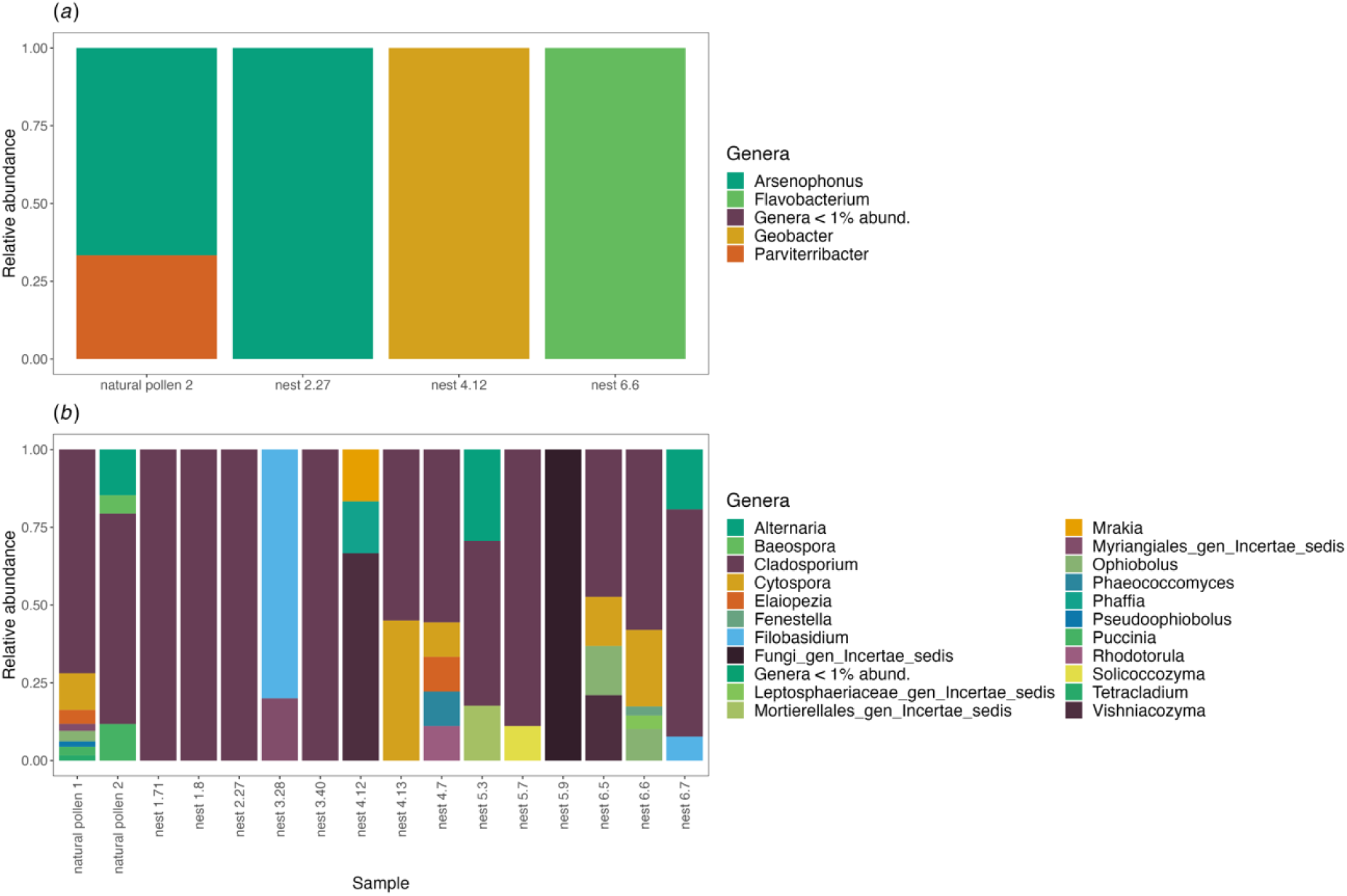
Relative abundances (proportion of sequence reads) of (*a*) bacterial and (*b*) fungal genera present in provisions before and after homogenization.

**Figure S3.**
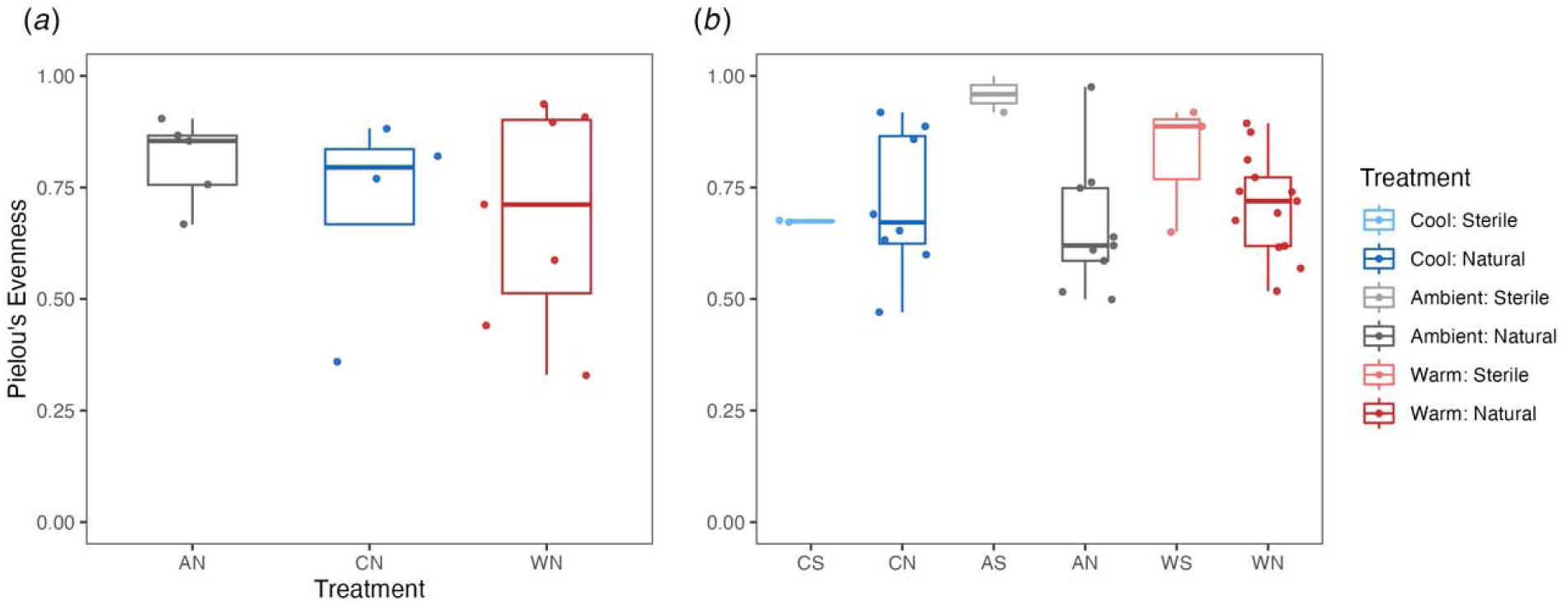
Pielou’s evenness of provisions with male bees using (*a*) bacterial and (*b*) fungal amplicon sequencing variants (ASVs). Temperature treatment and microbiome treatment were not significant predictors in neither the bacterial nor fungal model.

**Figure S4.**
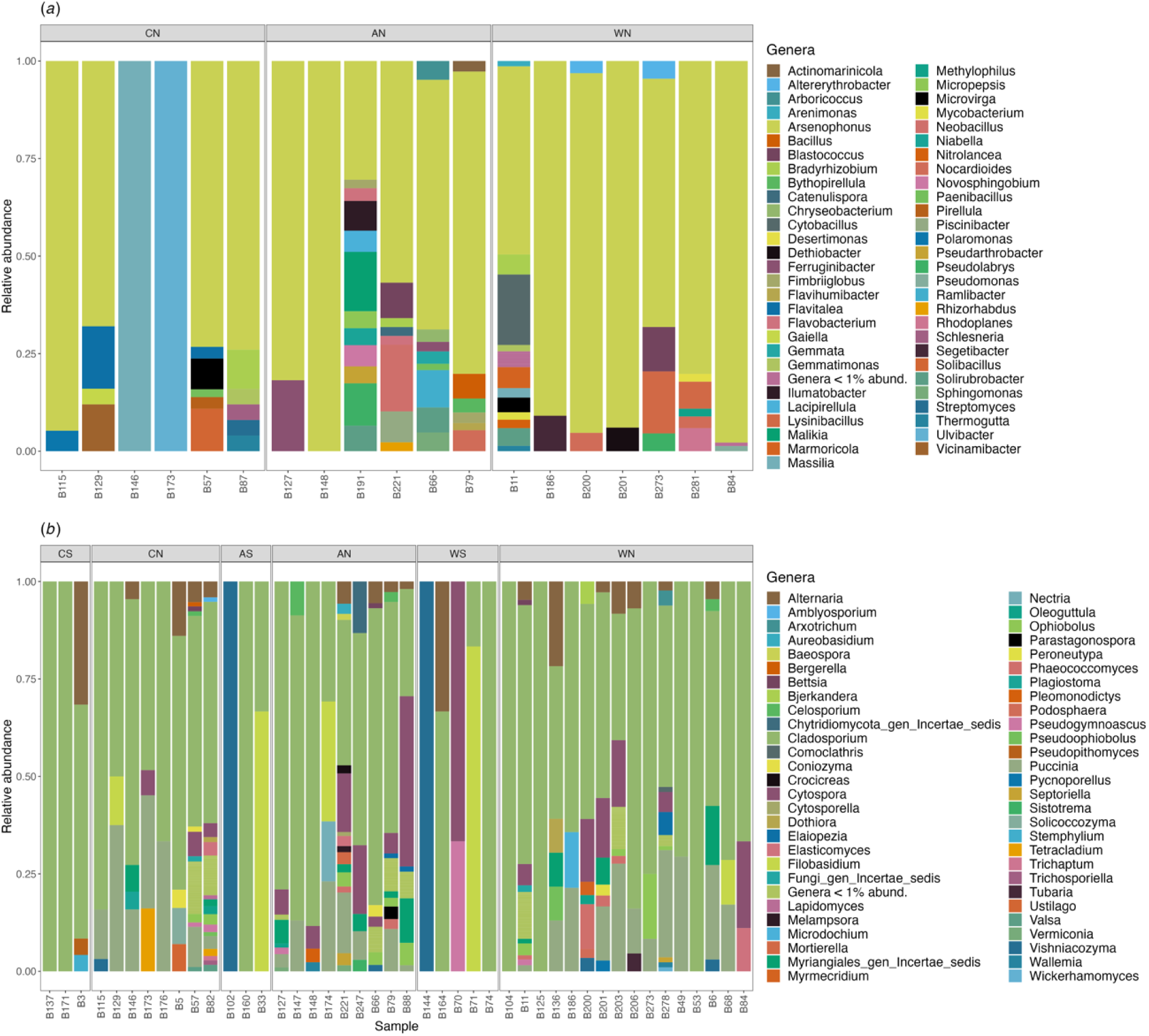
Relative abundance (proportion of sequence reads) of (*a*) bacterial and (*b*) fungal genera present in provisions with bees.

**Figure S5.**
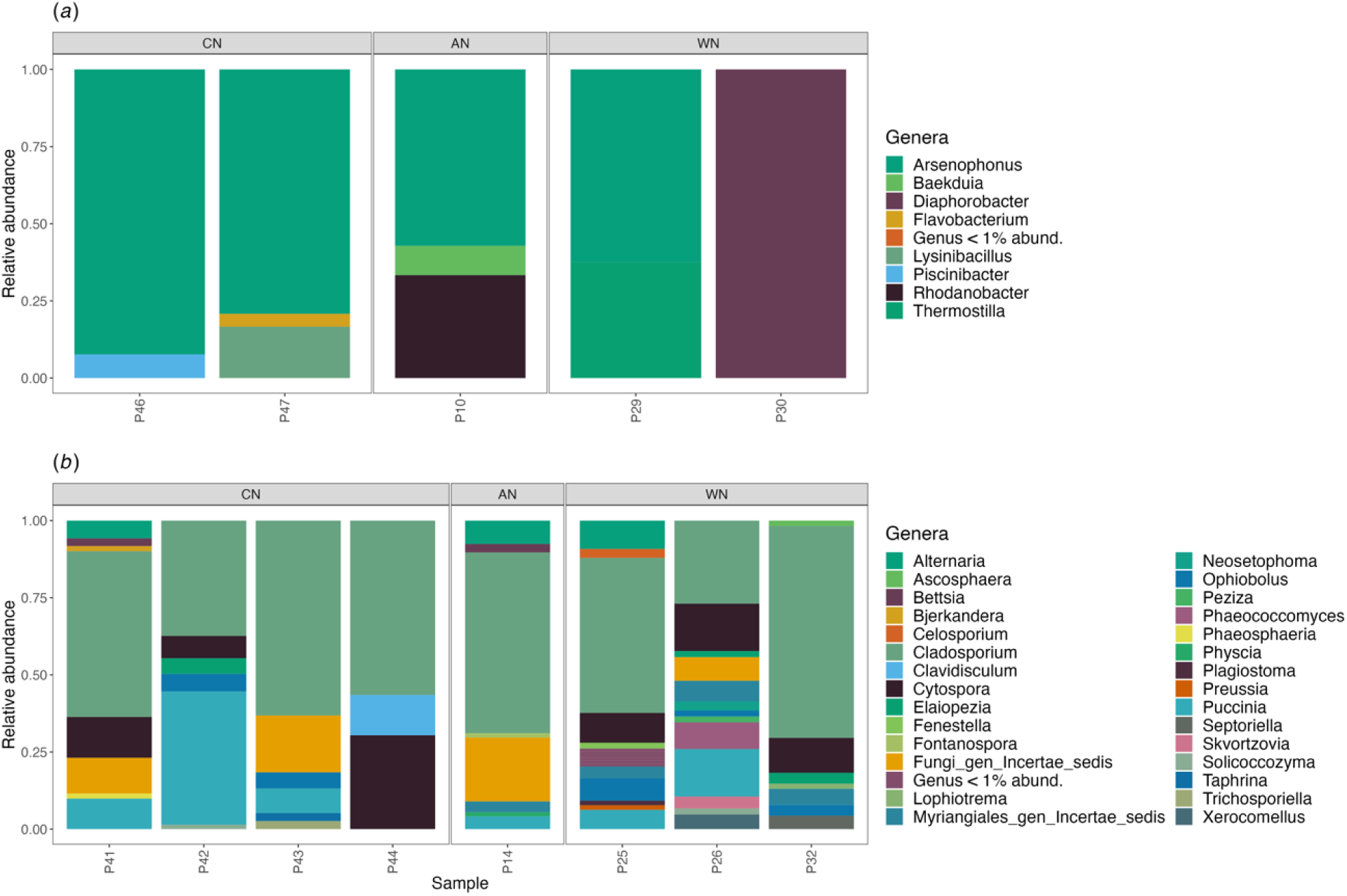
Relative abundance (proportion of sequence reads) of (*a*) bacterial and (*b*) fungal genera present in provisions without bees.

**Figure S6.**
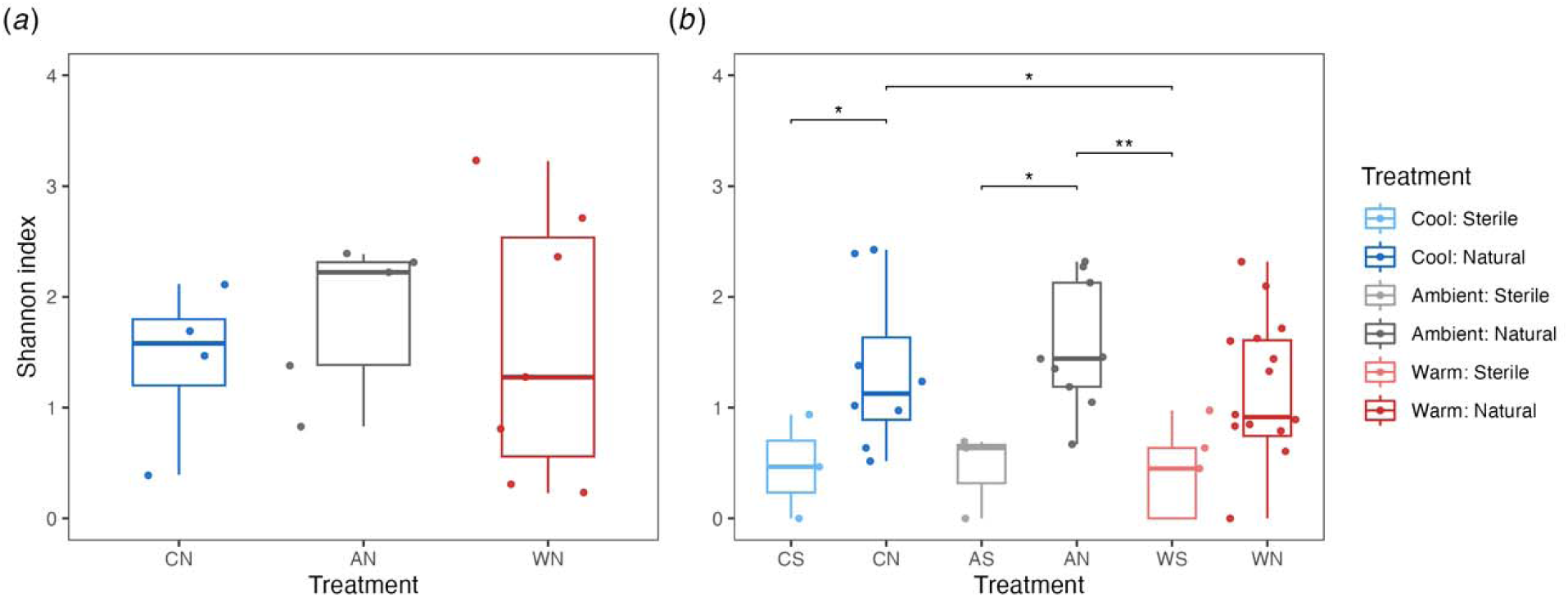
Shannon diversity indices of provisions with male and female bees using (*a*) bacterial and (*b*) fungal amplicon sequencing variants (ASVs). Temperature treatment was not a significant predictor in the bacterial model. However, microbiome treatment was a significant covariate (*p* = 0.0001) in the fungal model. Asterisks represent significant differences, where ‘*’ indicates a *p*-value < 0.05, ‘**’ *p* < 0.01, and ‘***’ *p* < 0.001.

**Figure S7.**
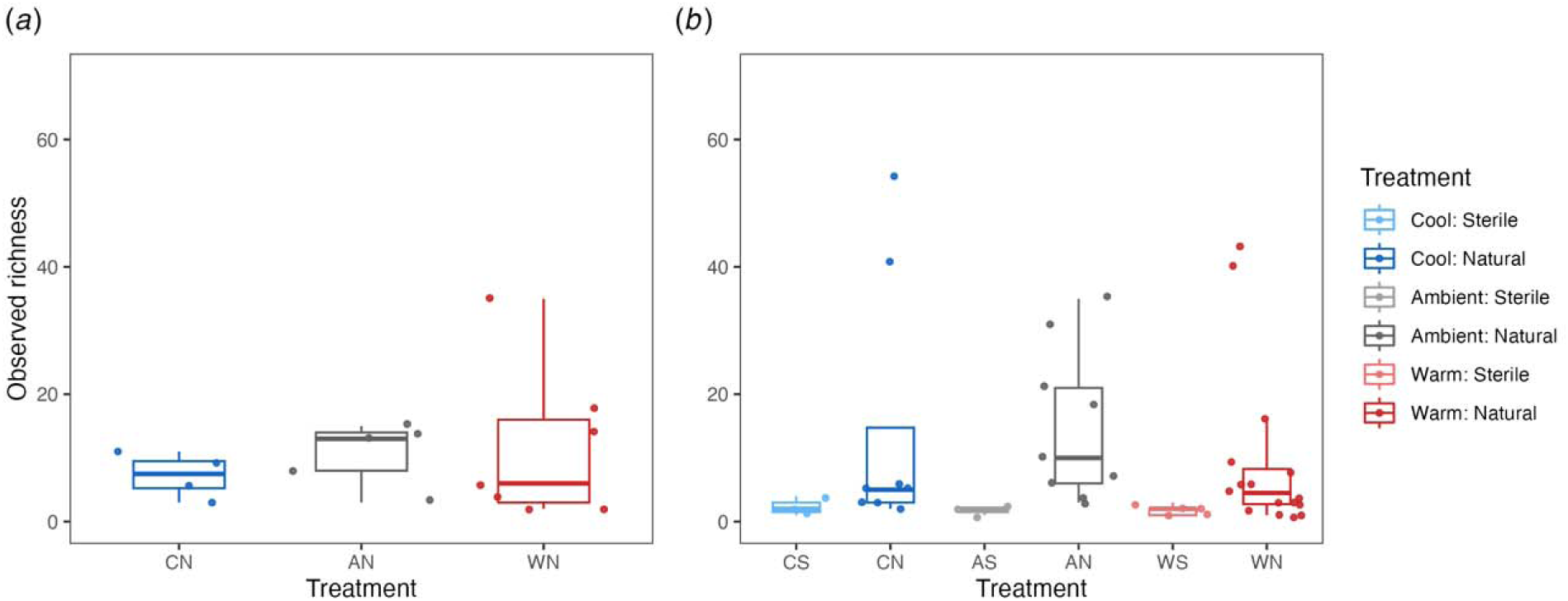
Observed richness of provisions with male and female bees using (*a*) bacterial and (*b*) fungal amplicon sequencing variants (ASVs). Temperature treatment was not a significant predictor (p = 0.66), yet bee sex had a significant effect on the observed richness of bacteria (p < 0.05). However, post-hoc tests did not reveal any significant pairwise comparisons. In the fungal model, both bee sex (p = 0.41) and microclimate (p = 0.42) did not affect observed richness. Rather, microbiome treatment had an effect on the observed richness of fungi (p < 0.05).

**Figure S8.**
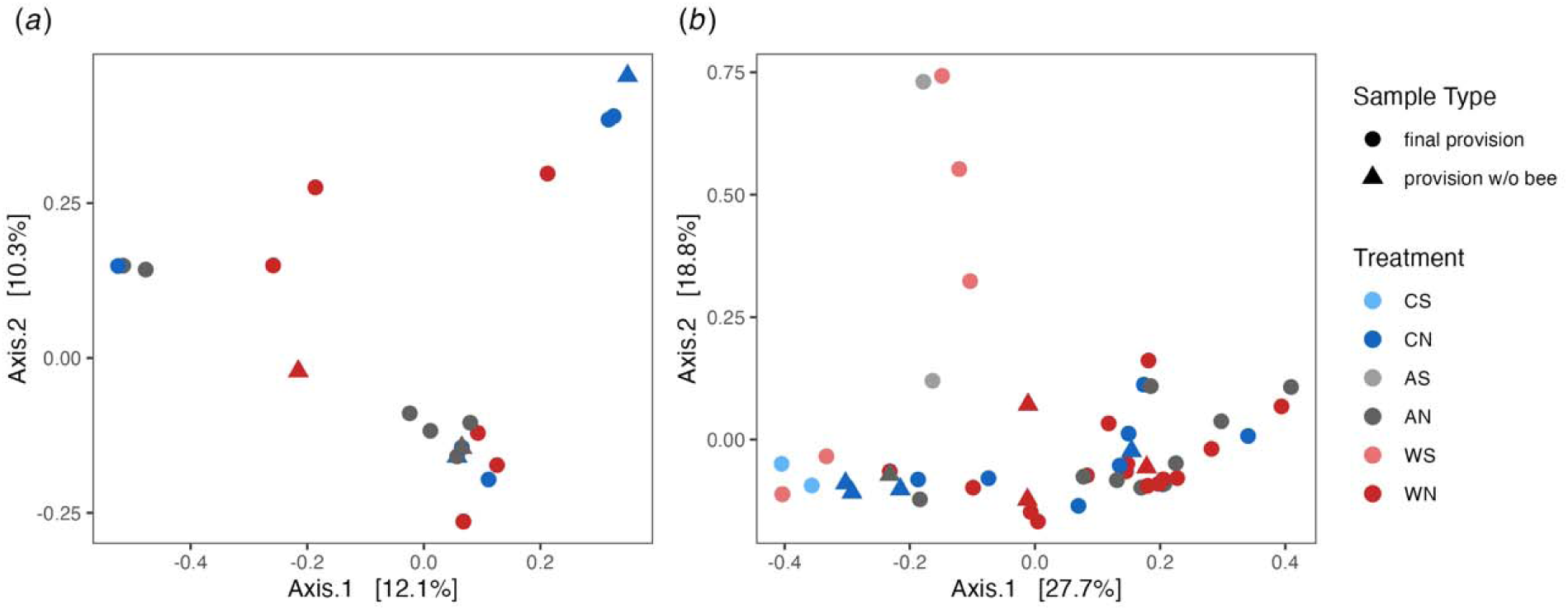
Principle coordinates analysis (PCoA) of Bray Curtis dissimilarity in provisions with and without bees of (*a*) bacterial and (*b*) fungal amplicon sequencing variants (ASVs). Bacterial composition was not significantly different. Microbiome treatment (*p* = 0.001) was a significant predictors of fungal composition.

**Table S1.** BLASTn results for *Arsenophonus* ASVs found with *Osmia lignaria* provisions. All 16 ASVs had high sequence similarity to the same accession (NR_042811.1), with a query cover of 100%. These ASVs accounted for a total of 835 bacterial reads (47.02% of total bacterial reads in all provision samples).

| ASV | Total Reads<br>per ASV | Percent Identity |
| --- | --- | --- |
| ASV1 | 120 | 98.79% |
| ASV2 | 95 | 98.79% |
| ASV3 | 87 | 99.03% |
| ASV4 | 74 | 98.79% |
| ASV5 | 68 | 98.79% |
| ASV6 | 66 | 99.02% |
| ASV7 | 59 | 99.51% |
| ASV8 | 57 | 98.54% |
| ASV9 | 36 | 99.03% |
| ASV10 | 34 | 99.27% |
| ASV11 | 32 | 99.27% |
| ASV12 | 31 | 99.03% |
| ASV13 | 28 | 99.03% |
| ASV14 | 27 | 98.79% |
| ASV15 | 12 | 98.30% |
| ASV16 | 9 | 98.79% |

